# racoon_clip – a complete pipeline for single-nucleotide analyses of iCLIP and eCLIP data

**DOI:** 10.1101/2024.02.27.582237

**Authors:** Melina Klostermann, Kathi Zarnack

## Abstract

**Summary:** Here, we introduce `racoon_clip`, a sustainable and fully automated pipeline for the complete processing of iCLIP and eCLIP data to extract RNA binding signal at single-nucleotide resolution. `racoon_clip` is easy to install and execute, with multiple pre-settings and fully customizable parameters, and outputs a conclusive summary report with visualizations and statistics for all analysis steps.

**Availability and Implementation:** `racoon_clip` is implemented as a snakemake-powered command line tool (snakemake version ≥ 7.22, Python version ≥ 3.9). The latest release can be downloaded from GitHub (https://github.com/ZarnackGroup/racoon_clip/tree/main) and installed via pip. A detailed documentation, including installation, usage and customization, can be found at https://racoon-clip.readthedocs.io/en/latest/.

The example datasets can be downloaded from the Short Read Archive (SRA; iCLIP: SRR5646576, SRR5646577, SRR5646578) or the ENCODE Project (eCLIP: ENCSR202BFN).

**Contact:** Kathi Zarnack

**Issue Section:** Genome analysis

## Introduction

UV crosslinking and immunoprecipitation (CLIP) coupled to high-throughput sequencing has become a popular tool to query the RNA binding behavior of RNA-binding proteins (RBPs) in a transcriptome-wide manner (Lee and Ule, 2018; Ule *et al*., 2003). CLIP protocols use UV irradiation to crosslink the RBPs to their bound RNAs *in vivo* and then extract the RBP-RNA complexes via immunoprecipitation with a specific antibody. Two common variants of CLIP, named individual-nucleotide resolution CLIP (iCLIP) (König *et al*., 2010) and enhanced CLIP (eCLIP) (Van Nostrand *et al*., 2016), detect the crosslink sites of RBPs on the RNAs by capturing the truncation of reverse transcription at these sites (**Figure 1A**). From these data, the crosslink position of the RBP can be extracted with single-nucleotide resolution as the crosslink sites reside one nucleotide (nt) upstream of the 5’ ends of the reads (Haberman *et al*., 2017). Still, there are differences between the protocols like the positioning and design of experimental barcodes and unique molecular identifiers (UMIs) or the preferred usage of single-end versus pair-end sequencing (**Figure 1B, Table 1**). Therefore, while the results of iCLIP and eCLIP experiments should be very comparable from an experimental point of view, differences in the experimental design and the resulting read architecture must be considered in the computational analysis (Buchbender *et al*., 2020).

**Table 1:** Differences of iCLIP and eCLIP data types that are relevant for data processing.

|  | **iCLIP**<br>(König *et al.*,<br>2010) | **iCLIP2**<br>(Buchbender *et al.*, 2020) | **eCLIP**<br>(Van Nostrand<br>*et al.*, 2016) | **eCLIP**<br>downloaded<br>from ENCODE |
| --- | --- | --- | --- | --- |
| **Library**<br>**strategy** | Single-end reads | Single-end reads | Paired-end reads<br>(read #2<br>contains<br>crosslink) | Paired-end reads<br>(read #2<br>contains<br>crosslink) |
| **UMI (N) and**<br>**barcode (X)**<br>**setup** | N<sub>3</sub>X<sub>4</sub>N<sub>2</sub> | N<sub>5</sub>X<sub>6</sub>N<sub>4</sub> | Read #2: N<sub>5</sub> or<br>N<sub>10</sub> | Read #2: N<sub>5</sub> or<br>N<sub>10</sub> already<br>removed from<br>sequence and<br>stored in read<br>name |

**Figure 1:**
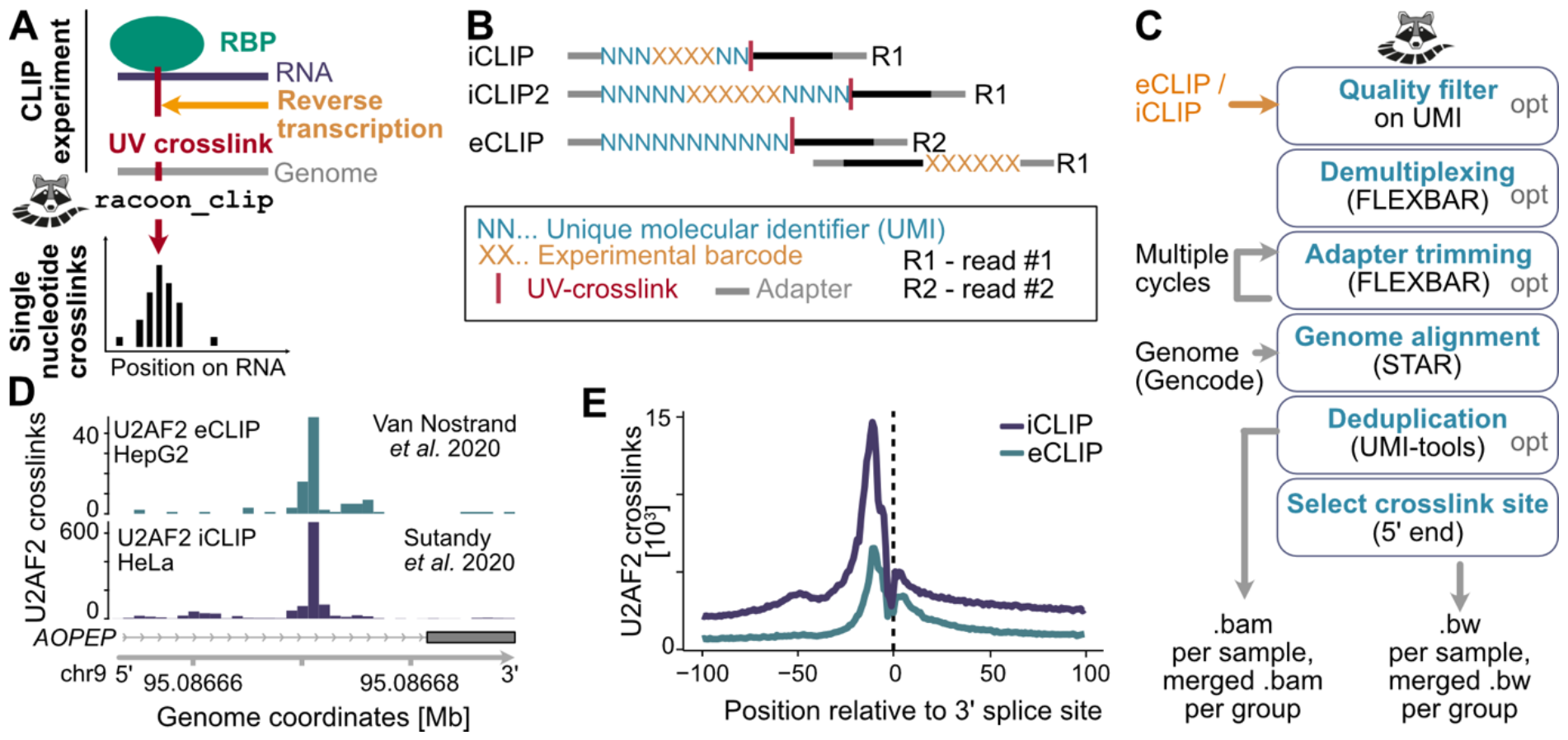
*racoon_clip* allows for efficient processing of iCLIP and eCLIP data. **(A)** Schematic of iCLIP/eCLIP experiments (left) to obtain single-nucleotide resolution (right). The RNA-binding protein (RPB, green) of interest is crosslinked to its *in vivo* RNA targets (dark blue) via UV irradiation. The RBP–RNA complex is immunoprecipitated with a specific antibody. Reverse transcription (orange arrow) of the RNA fragment will stop one nucleotide upstream of the UV crosslink site (red bar). By extracting this position, iCLIP/eCLIP data can be processed to single-nucleotide resolution. **(B)** Differences in the sequencing read composition between iCLIP, iCLIP2 and eCLIP experiments. All experiments carry the unique molecular identifier (UMI) at the 5’ end of the truncated cDNAs. However, since eCLIP usually employs paired-end sequencing, read #2 will contain the truncation site. The experimental barcode in iCLIP and iCLIP2 is positioned between the UMIs, while it is separately positioned in read #1 in eCLIP. **(C)** Steps performed by `racoon_clip`. **(D)** Exemplary crosslink profiles of U2AF2 eCLIP (Van Nostrand, Pratt, *et al*., 2020) (top) and U2AF2 iCLIP (Sutandy *et al*., 2018) (bottom) after `racoon_clip` processing. **(E)** Metaprofile of U2AF2 crosslink events from eCLIP (petrol) and iCLIP (dark blue) data around annotated 3’ splice sites.

Here, we present the command line tool `racoon_clip` that implements single-nucleotide resolution processing for iCLIP and eCLIP data. The automated pipeline is based on the previously published workflow (Busch *et al*., 2020) that was extended to cater both iCLIP and eCLIP data. Being built from a snakemake pipeline (Mölder *et al*., 2021; Roach *et al*., 2022), `racoon_clip` can be run in a fully automated and multi-threaded manner. Furthermore, it can be easily installed via download from GitHub and local installation via pip. Together, this makes `racoon_clip` an easy-to-use option to obtain single-nucleotide information on RBP crosslink sites from iCLIP and eCLIP data.

### Workflow

`racoon_clip` processes raw iCLIP or eCLIP sequencing reads to obtain single-nucleotide RBP crosslink sites. It is a command line tool powered by a snakemake pipeline implemented in Python, bash and R and uses publicly available command line tools where possible. `racoon_clip` can be downloaded from GitHub (https://racoon-clip.readthedocs.io/en/latest/) and installed via pip, including all dependencies.

`racoon_clip` can be run on data from different CLIP protocols by either specifying one of the pre-set experiment types (“iCLIP”, “iCLIP2”, “eCLIP_5ntUMI”, “eCLIP_10ntUMI”, “eCLIP_ENCODE_5ntUMI”, “eCLIP_ENCODE_10ntUMI”, or “noBarcode_noUMI”) with the *experiment_type* parameter or using a custom barcode and UMI setup. As input, `racoon_clip` takes the sequencing reads as multiplexed or demultiplexed FASTQ files, the genome assembly as FASTA file, and the gene annotation as GTF file. All steps performed by `racoon_clip` are fully customizable and can be specified either via a configuration (config) file or directly in the command line (see User manual). The output of `racoon_clip` includes the aligned reads in BAM format, the single-nucleotide crosslink events in BED and BIGWIG format, as well as a detailed processing report in HTML format.

### Performed steps

The `racoon_clip` pipeline consists of three major steps (**Figure 1C**): (1) the preprocessing of the sequencing reads, (2) the genomic alignment, and (3) the extraction of crosslink events. In the preprocessing step, `racoon_clip` deals with the barcodes of the iCLIP or eCLIP experiment, taking into account multiple barcode formats. In brief, the FASTQ files are demultiplexed, the barcodes are trimmed off, and the UMIs are stored in the read names. To perform this on data from different protocols, `racoon_clip` offers predesigned settings for iCLIP, iCLIP2, eCLIP or eCLIP downloaded from ENCODE and a flexible architecture of experimental barcodes with the basic structure U_UMI1_len_–B_barcode_len_–V_UMI2_len_ for custom read designs.

`racoon_clip` can perform barcode and/or adapter trimming and/or demultiplexing using FLEXBAR (Dodt *et al*., 2012) (version 3.5.0). These options can be specified depending on the input data. During barcode trimming, the UMIs are appended to the read names for later deduplication. If present, barcodes need to be provided in a FASTA file. To ensure correct barcode and UMI assignment, `racoon_clip` can perform an optional quality filtering on the barcode regions (*quality_filter_barcodes*) using awk (mawk version 1.3.3). Of note, there is a specialized option for eCLIP data directly downloaded from the ENCODE website, as ENCODE provides preprocessed sequencing reads without barcodes and with UMIs already stored in the read name, albeit in a different position. For adapter trimming, custom adapters can be provided as a FASTA file. Otherwise, a standard set of sequencing adapters from Illumina and eCLIP adapters is used. By default, one round of adapter trimming is performed but multiple cycles can be chosen. For example, two cycles are recommended for ENCODE eCLIP data (https://github.com/yeolab/eclip). We advice to check in the HTML report that adapters have been trimmed off correctly. The output of the preprocessing step are demultiplexed FASTQ files, containing the sequencing reads of the individual samples without adapters and barcodes and with the UMIs appended to the read names.

In the second step, the demultiplexed reads are aligned to the genome and then deduplicated. For genomic alignment using STAR (Dobin *et al*., 2013) (version 2.7.10), the FASTA file of the genome assembly and the corresponding GTF file of the gene annotation need to be provided. Soft-clipping is restricted to the 3’ end of the reads (*--alignEndsType* “Extend5pOfRead1”) to preserve the exact crosslink position upstream of the 5’ end. All other STAR parameters can be customized, for instance to allow for multi-mapping reads. By default, STAR is configured to output only unique alignments (--*outFilterMultimapNmax* 1) with up to 4% mismatches (--*outFilterMismatchNoverReadLmax* 0.04). Next, the aligned reads are deduplicated by removing reads mapping at the same location and sharing the same UMI using UMI-tools (Smith *et al*., 2017) (version 1.1.1). There is an option to skip deduplication (*deduplicate: False*), although we recommend performing deduplication if possible. Finally, the BAM files are indexed using SAMtools (Li *et al*., 2009) (version 1.11). Ultimately, the second step returns aligned, deduplicated reads in BAM format.

In the third step of `racoon_clip`, the crosslink sites are extracted by selecting the position 1 nt upstream of the 5’ end of the aligned reads using BEDTools (Quinlan and Hall, 2010) (version 2.30.0) and kentUtils (https://github.com/ucscGenomeBrowser/kent) (version 377).

The crosslink events are output in BED and BIGWIG format.

As it is often useful to have merged files of samples from the same condition, sample groups can be specified. In this case, `racoon_clip` provides an additional merged file for each sample group. If no group is specified, all samples will be merged by default.

Once completed, `racoon_clip` provides an HTML processing report that summarizes the data quality (using FastQC (Andrews, 2010) (version 0.12) and MultiQC (Ewels *et al*., 2016) (version 1.14)) and relevant statistics for each step. This allows the user to assess the quality of the processed dataset and ensure that all steps were performed as expected.

## Application

To showcase the functionalities of `racoon_clip`, we processed an eCLIP dataset from ENCODE (Van Nostrand, Pratt, *et al*., 2020; Van Nostrand, Freese, *et al*., 2020) (2 replicates, 18,760,220 reads in total; ENCODE ID: ENCSR202BFN) and an iCLIP dataset from a previous publication (Sutandy *et al*., 2018) (3 replicates, 66,041,912 reads in total; SRA ID: SRR5646576, SRR5646577, SRR5646578) for the splicing factor U2AF2. For the iCLIP dataset, `racoon_clip` needed 1 hour and 54 minutes at a peak RAM usage of 40.5 Gigabytes on 6 cores. As the data is stored at SRA already demultiplexed, deduplicated and with adapters and barcodes trimmed, we used the following settings for the processing: *experiment_type: iCLIP, demultiplex: False, quality_filter_barcodes: False, adapter_trimming: False, deduplicate: False*.

For the eCLIP dataset, 2 hours and 13 minutes at a peak RAM usage of 35.4 Gigabytes were needed on 6 cores. The following settings were used: *experiment_type: eCLIP_ENCODE_10ntUMI, demultiplex: False, quality_filter_barcodes: False, adapter_trimming: True, adapter_cycles: 2, deduplicate: True*.

From the iCLIP data, 95.8% of the sequencing reads were uniquely aligned. In total, `racoon_clip` detected 63,266,600 crosslink events for the merged replicates. Similarly, for the eCLIP data, 67.2% were uniquely mapped, and of these 81.2% were kept after deduplication, yielding a total of 10,282,489 crosslink events.

Exemplary U2AF2 crosslink profiles from the iCLIP and eCLIP data on the *AOPEP* transcript are shown in **Figure 1D**. The unified processing by `racoon_clip` makes both datasets directly comparable. As expected, the U2AF2 crosslink events accumulate upstream of 3’ splice sites (**Figure 1E**), underlining the high resolution of the RBP crosslink profiles.

## Conclusion

In summary, `racoon_clip` enables users to obtain single-nucleotide resolution signal from their eCLIP and iCLIP data. The output files can directly be used as input for binding site definition as described in (Busch *et al*., 2020), using for example the peak calling software PureCLIP (Krakau *et al*., 2017) and R/Bioconductor package BindingSiteFinder (Brüggemann and Zarnack). `racoon_clip` thereby builds a solid foundation for in-depth eCLIP and iCLIP data analyses and facilitates comparisons of multiple experiments from rich resources such as the ENCODE eCLIP data (Luo *et al*., 2019; Van Nostrand, Pratt, *et al*., 2020).

## Acknowledgements

We thank You Zhou and Dominik Stroh and testing `racoon_clip` and its documentation and the Zarnack group for help and discussions.

